# STAT3 is a genetic modifier of TGF-beta induced EMT in KRAS mutant pancreatic cancer

**DOI:** 10.1101/2023.09.01.555946

**Authors:** Stephen D’Amico, Varvara Kirillov, Oleksi Petrenko, Nancy C. Reich

## Abstract

Oncogenic mutations in KRAS are among the most common in cancer. Classical models suggest that loss of epithelial characteristics and the acquisition of mesenchymal traits are associated with cancer aggressiveness and therapy resistance. However, the mechanistic link between these phenotypes and mutant KRAS biology remains to be established. Here we identify STAT3 as a genetic modifier of TGF-beta-induced epithelial to mesenchymal transition. Gene expression profiling of pancreatic cancer cells identifies more than 200 genes commonly regulated by STAT3 and oncogenic KRAS. Functional classification of STAT3 responsive program reveals its major role in tumor maintenance and epithelial homeostasis. The signatures of STAT3-activated cell states can be projected onto human KRAS mutant tumors, suggesting that they faithfully reflect characteristics of human disease. These observations have implications for therapeutic intervention and tumor aggressiveness.

**Significance:** The identification of the molecular and genetic characteristics of tumors is essential for understanding disease progression and aggressiveness. KRAS mutations are the most frequent oncogenic drivers in human cancer. In this study we demonstrate that the ubiquitously expressed transcription factor STAT3 is a genetic modifier of TGF-beta-induced EMT, and thereby oncogenic KRAS dependency. Both *in vitro* and *in vivo* studies demonstrate that STAT3 responsive program is an inherent part of oncogenic KRAS outcome.

## Introduction

Pan-cancer projects, such as The Cancer Genome Atlas (TCGA), have provided a comprehensive view of the mutational landscape in human cancers. The foremost objective has been the discovery of key genes that drive cancer initiation and progression. It is estimated that cancer genomes contain an average of less than five driver mutations, whose outcomes are realized in the context of chromosomal and epigenetic alterations (1–3). While the Cancer Gene Census (CGC) has been largely defined, unraveling the contributions of normal cell functions to cancer development and their influence on tissue homeostasis, plasticity, and differentiation remains a complex task. A large body of evidence suggests that signal transducer and activator of transcription 3 (STAT3) has tumor-promoting properties that it exerts in a context-dependent fashion (4, 5). Canonical activation of STAT3 occurs following phosphorylation of tyrosine 705 (pY705) by receptor-associated Janus kinases (JAKs) or other tyrosine kinases (6). The clinical relevance of hyperactive STAT3 has been linked to subsets of hematological malignancies, with the identification of JAK1/3 or STAT3 mutations (7–10). The most common STAT3 mutations, Y640F and D661Y, render STAT3 constitutively active (7, 8). In sharp contrast, STAT3 mutations rarely occur in solid tumors. TCGA pan-cancer analysis reveals that most cancers do not express high levels of activated STAT3 (www.cancer.gov). Patient-derived xenografts and genetically engineered mouse models have yielded contrasting findings regarding the role of STAT3 in cancer development that range from tumor-promoting to tumor-suppressive, suggesting a high degree of tissue specificity (5).

We aim to delineate the role of STAT3 in shaping the patterns of oncogenic KRAS dependency in KRAS mutant cancer cells. A previous study from our laboratory uncovered a novel link between STAT3 and cancer showing that activation of STAT3 in KRAS mutant cancers led to the stabilization of epithelial differentiation (11). This observation suggests that STAT3 plays a dynamic role in in modulating the phenotypic diversity of KRAS-driven tumors, ostensibly coupled with the selection of the fittest. In this study we leverage isogenic STAT3 intact and deficient cells to more fully delineate the effects of STAT3 on oncogenic KRAS dependency and the growth of cancer cells in culture or as tumors. To determine whether KRAS dependent tumor cells are co-dependent on STAT3, we used two well-established models of KRAS mutant cancer: mouse embryonic fibroblasts and pancreatic ductal adenocarcinoma (PDAC) cells expressing endogenous KRAS^G12D^. Both *in vitro* and *in vivo* assays demonstrate that neither persistent activation of STAT3 nor its loss confers distinct growth advantages on tumor cells. Instead, STAT3 guides morphological and functional characteristics of the transformed cells and tumors. Stabilization of the epithelial phenotype and attenuation of the TGF-β/SMAD4 pathway are two main driving forces behind STAT3 activity (12–15). The data highlight antagonistic epistasis between SMAD4 and STAT3, where SMAD4 expressing tumors are poorly differentiated and exhibit mesenchymal features only in the absence of STAT3, while SMAD4 deficient tumors are well-differentiated and display epithelial morphology only in presence of STAT3. The results have implications for our understanding of the molecular basis of oncogenic KRAS dependency and therapy response.

## Materials and Methods

### Cells and Reagents

Clonally-derived KRAS^G12D^ p53KO mouse embryonic fibroblasts (KP MEFs) and pancreatic KRAS^G12D^ p53^R172H^ (KPC) cells were previously described (16, 17). Human Hep3B and HEK293T cells were obtained from ATCC. All cells were grown in DMEM media supplemented with 5% FBS (Atlanta Biologicals) and 1x antibiotic/antimycotic (Gibco). For standard proliferation assays, cells were seeded into 6-well plates at a concentration of 4×10^5^ cells per well and counted cumulatively with a Coulter counter (Beckman) every three days for two weeks. Focus formation assays were performed as described (16, 18). Briefly, 10^3^ KP MEFs were co-cultured with 10^5^ p53KO feeder MEFs in 6 cm dishes. After two weeks, transformation efficiency was evaluated by manually counting macroscopic colonies. Transformed foci were stained with Giemsa (Sigma) for visualization. Stable KP MEF and KPC knockout cell lines were generated via selection in 2 μg/ml puromycin followed by single colony isolation. In vitro luciferase activity was measured using a Lumat model LB luminometer (Promega) and the Luciferase Reporter Gene Assay according to manufacturer’s instructions (Roche). Hep3B cells were stimulated with human IL-6 (Cell Signaling Technology) at a concentration of 20ng/ml for 24 hours.

### Lentivirus and plasmids

Lentiviral (adapted from the pWPXL/pEF1a backbone) and pEGFP-N1 (Addgene) expression vectors encoding mutant STAT3 alleles were derived using site-directed mutagenesis. The final plasmids were sequence confirmed. The p3XGAS-Hsp70-Luc reporter plasmid was described (19). For CRISPR/Cas9-mediated knockouts, we used single guide RNAs (sgRNAs) for STAT3 (5’-gcagctggacacacgctacc-3’ or 5’-gtacagcgacagcttcccca-3’), SMAD4 (5′-ggtggcgttagactctgccg-3′), and TGFBR2 (5’-ccttgtagacctcggcgaag-3’) cloned into LentiCRISPRv2 puro (Addgene) (20). Lentiviruses were produced by transient transfection of HEK293T and collected according to standard protocols.

### Tumorigenicity in Mice

All animal studies were approved by the Institutional Animal Care and Use Committee at Stony Brook University. Male NU/J (nude) mice (5 weeks old) (The Jackson Laboratory) were inoculated orthotopically or subcutaneously with 10^4^ cells in 100 μl of Matrigel (Corning) diluted 1:7 with Opti-MEM (Corning). Orthotopic implantations into the pancreas were performed using standard procedures (21). Pancreatic tumor latency was determined through abdominal palpation. We defined subcutaneous tumor latencies as the period between implantation of tumorigenic cells into mice and the appearance of tumors 1 mm in diameter. The end point was a tumor diameter of 0.5 cm. Statistical analyses were performed using two-tailed Student’s t-test at the 95% confidence interval. P ≤ 0.05 was considered statistically significant. Tumor-initiating cell (TIC) frequency was determined by extreme limiting dilution assays and online ELDA software (http://bioinf.wehi.edu.au/software/elda/). The number of cells in each subcutaneous injection ranged from 10^2^ to 10^4^. Mouse tumor tissue was harvested, fixed in 5 volumes of 4% paraformaldehyde for 48 hours, and processed via the Stony Brook University Histology Core. Paraffin-embedded formalin-fixed 5 μm sections were stained with hematoxylin and eosin for histology.

### Expression analysis

Western blotting was performed using antibodies against AKT1 (4691), P-AKT1 pS473 (4060), CDH1 (3195), P-ERK1/2 (4370), SMAD4 (46535), P-STAT3 pY705 (9131), VIM (5741) (all from Cell Signaling), ERK1/2 (05-157, Millipore), STAT3 (610190, BD) and TGFBR2 (sc-400, Santa Cruz). Whole cell extracts were prepared by lysing cells in buffer containing 10 mM Tris HCl, pH7.4, 150 mM NaCl, 1 mM EDTA, 10% glycerol, 1% Triton X100, 40 mM NaVO4, 0.1% SDS, and 1x protease inhibitors (Roche). Western blots were imaged using Image Studio software (LI-COR). Total cellular RNA was isolated using PureLink RNA (ThermoFisher) according to manufacturer’s specifications and phenol-extracted. Pancreatic tissues were incubated for 24 hours at 4°C in ≥5 volumes of RNAlater solution (ThermoFisher) to preserve RNA integrity. RNA sequencing and bioinformatics were performed by Novogene Corporation (http://en.novogene.com). STAT3 and SMAD4 knockout signature scores (STAT3KO_UP, STAT3KO_DN, SMAD4KO_UP, SMAD4KO_DN) were computed as the average of RNA expression values (fpkm) of the top up- or down-regulated genes in KPC cell lines. EMT and mouse Ras Dependency Index (RDI) scores were calculated using defined gene sets (22–24). For publicly available human datasets, RDI, KRAS_sig, RSK_sig, epithelial (EPI) and mesenchymal (MES) gene expression scores were calculated as the sum of RNA expression values (z-scores) using previously characterized gene modules (25, 26).

### Statistics and Reproducibility

Statistical analysis was performed using two-tailed Student’s t-test, Fisher’s exact test or Wilcoxon test, as appropriate for the dataset. An FDR adjusted p-value (q-value) was calculated for multiple comparison correction. Individual mice and tumor cell lines were considered biological replicates. Statistical details for each experiment are denoted in the corresponding figures and figure legends. The micrographs (H&E) represent at least three independent experiments. All data are presented as mean ± SD. In box and whisker plots, the middle line is plotted at the median, the upper and lower hinges correspond to the first and third quartiles, and the ends of the whisker are set at 1.5 x IQR above the third quartile and 1.5 x IQR below the first quartile (IQR, interquartile range or difference between the 25th and 75th percentiles).

## Results

### Effect of STAT3 activity on KRAS-mediated transformation

To assess the role of STAT3 in KRAS-driven tumorigenesis, we measured proliferation rates, contact inhibition, and tumor formation in mice. We have reported that p53-null mouse embryonic fibroblasts expressing endogenous mutant KRAS^G12D^ (termed KP MEFs) exhibit typical features of oncogenic transformation using quantitative and sensitive assays (16). CRISPR/Cas9-mediated gene editing was used to generate isogenic STAT3 knockouts in the KP MEFs, and gain of function (GOF) and loss of function (LOF) mutant STAT3 alleles were stably integrated into cells via lentiviral vectors (Figs. 1A, 1B). We used naturally occurring GOF mutants, Y640F, K658Y and D661Y, and a synthetic mutant STAT3C (A662C/N664C), which all render a persistently phosphorylated and thus hyperactive STAT3 pY705 (9, 27). LOF mutations targeted the STAT3 DNA binding (EE434-435AA and VVV461-463AAA) and transactivation domains (Y705F and S727A) (28) (Fig. S1A). All mutants exhibited relatively uniform expression levels that were ∼5-fold higher than endogenous STAT3 (Figs. 1B, S1B). As reported, STAT3 Y640F, K658Y and STAT3C displayed increased levels of Y705 phosphorylation. As neither STAT3 WT nor STAT3 GOFs exhibited robust phosphorylation on S727, we used a validated STAT3 mutant S727E that mimics the phosphorylation of S727 (29) (Fig. S1A).

**Figure 1.**
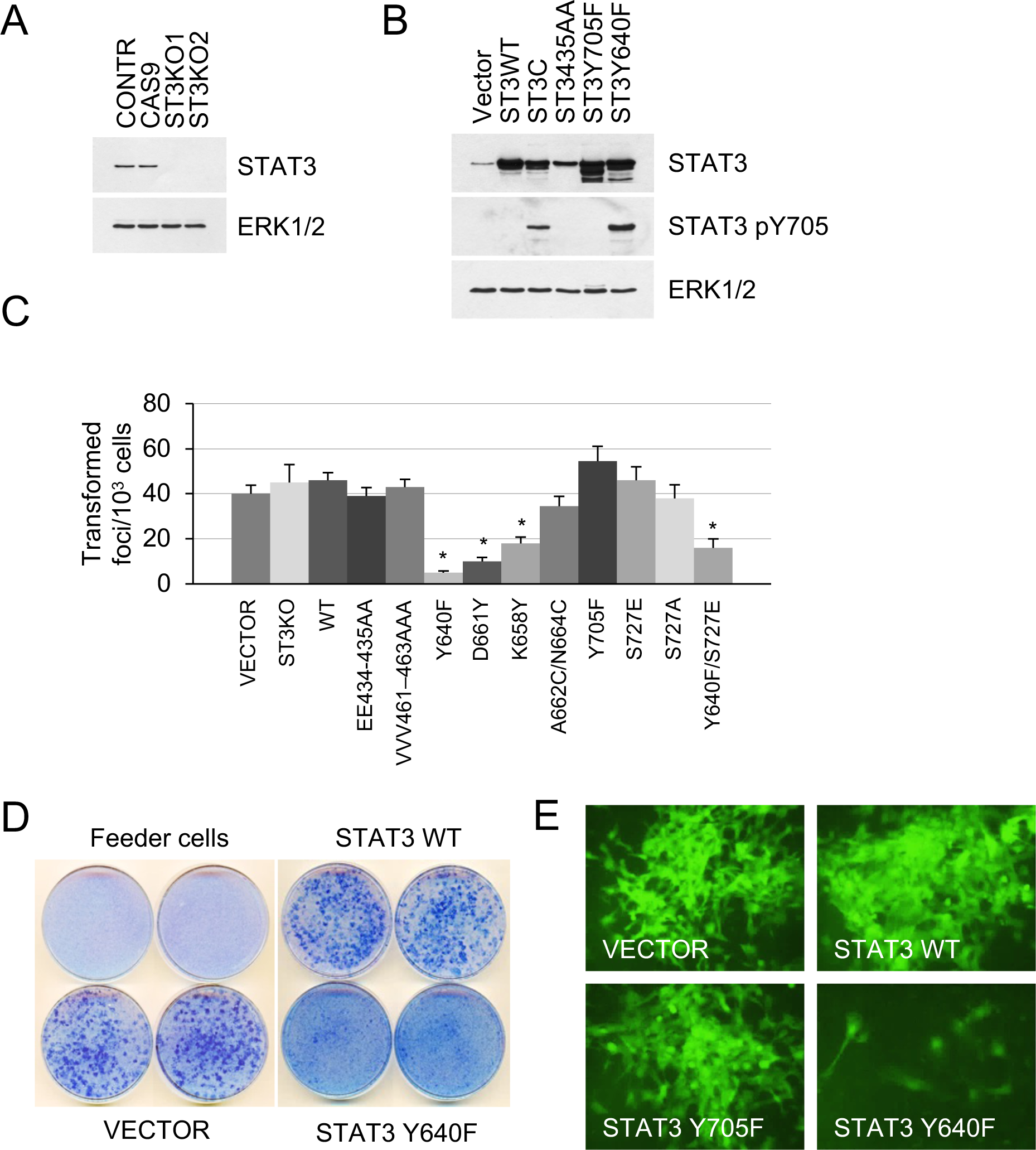
Effect of STAT3 activity on KRAS-mediated transformation. **A.** Western blot analysis of STAT3 expression in KP MEFs (Contr) transduced with CRISPR/CAS9 lentivirus expressing CAS9 or gRNAs targeting STAT3 (ST3KO). Two independent knockout clones are shown. ERK1/2 is a loading control. **B.** Western blot analysis of total and Y705-phosphorylated STAT3 in KP MEFs transduced with lentivirus expressing control (Vector), wild-type (ST3WT) or mutant STAT3 alleles as noted. ERK1/2 is a loading control. **C.** Transduced control (Vector) and KP MEFs were evaluated for foci formation. Cells expressing the indicated STAT3 genotypes were co-cultured with 10^3^ p53KO feeder MEFs and macroscopic colonies were counted after two weeks (n=3 for each cell type, *p<0.05). Values correspond to mean ± s.d. **D.** Representative images of tissue culture plates stained with Giemsa to detect foci formed by KP MEFs expressing vector, STAT3 WT or hyperactive STAT3 Y640F. **E.** Representative images of transformed foci visualized by fluorescence microscopy. Foci formed by KP MEFs co-expressing GFP with vector alone, STAT3 WT, STAT3 Y705F, or STAT3 Y640F are shown.

Isogenic KP MEF cell lines harboring wild-type or mutant STAT3 were evaluated for growth, tumorigenesis, and pathway activation. Loss of STAT3 expression did not affect cell growth under standard culture conditions (Fig. S1C). Likewise, loss of STAT3 did not affect KRAS-induced transformation. This is indicated by the ability of STAT3 KO cells to grow in multilayers and form transformed foci to the same extent as controls (Fig. 1C). Similar results were obtained using LOF mutations in STAT3 DNA binding (EE434-435AA and VVV461-463AAA) and transactivation domains (Y705F and S727A). In contrast, STAT3 GOF mutations, Y640F, D661Y, and K658Y, impaired KRAS-induced focus formation (p<0.005 by two-tailed T test, Figs. 1C, 1D). Because the transduced STAT3 constructs co-expressed a GFP reporter, the formation of transformed foci was visualized by fluorescence microscopy (Fig. 1E). Through these real-time studies we discovered that only a small percentage of cells with hyperactive STAT3 Y640F had some ability to form transformed colonies, while the majority of cells remained contact inhibited. We noted that STAT3 S727E had no significant effect on cell transformation, while the Y640F/S727E double mutant displayed only a marginal increase in the number of transformed foci relative to STAT3 Y640F itself (Fig. 1C). Thus, the hyperactive Y640F mutation exerts a dominant influence over S727E in KRAS-transformed MEFs. As expected, lentiviral expression of wild-type or GOF STAT3 alleles, Y640F and D661Y, failed to transform primary p53KO MEFs or immortalized NIH 3T3 cells (Fig. S1D, data not shown), indicating that STAT3 does not display intrinsic oncogenicity of its own.

### STAT3 GOF mutations reduce tumor development in mice

To investigate tumorigenic effects of STAT3 in vivo, subcutaneous implants of KP MEFs into nude mice were used. Tumors developed by STAT3 KO cells showed growth characteristics similar to those of STAT3 intact vector controls (Fig. 2A). In contrast, cells expressing the hyperactive STAT3 Y640F and, to a lesser extent, K658Y mutations were delayed in their ability to form tumors in mice. We used fluorescence-activated cell sorting for GFP to fractionate STAT3 Y640F MEFs into pools with low and high STAT3 pY705 expression (Fig. 2B). Following implantation in mice, cell populations with high STAT3 pY705 expression developed tumors more slowly compared to low expressing cells (p=0.005, Fig. 2C). Therefore, there is a dose-dependent ability of hyperactive STAT3 Y640F to limit tumorigenicity of KRAS-transformed MEFs. Limiting dilution assays in nude mice revealed that the frequency of tumor-initiating cells was reduced by approximately 9-fold in high STAT3 Y640F expressing cells compared to control cells (Fig. 2D).

**Figure 2.**
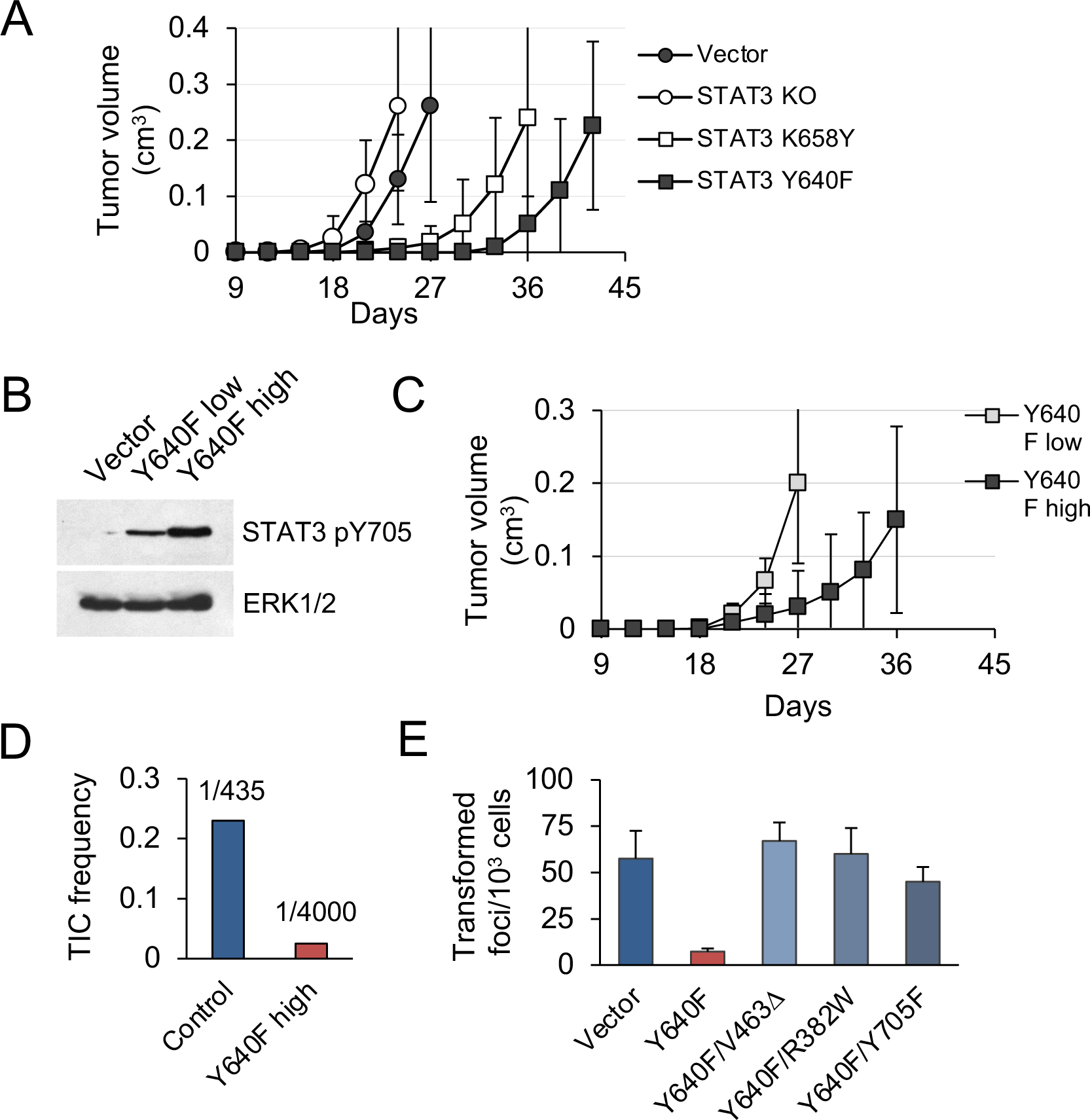
STAT3 GOF mutations reduce tumor development in mice. **A.** Subcutaneous tumor formation in nude mice by KP MEFs expressing vector alone, STAT3 KO, STAT3 K658Y or STAT3 Y640F (10^4^ cells per injection site, n=4 for each cell type). Error bars represent s.d. **B.** Western blot analysis of Y705-phosphorylated STAT3 in KP MEFs expressing vector alone, and low or high levels of STAT3 Y640F protein. Cells were fractionated by FACS for GFP. ERK1/2 is a loading control. **C.** Subcutaneous tumor formation in nude mice by cells from (B) (10^4^ cells per implant site, n=4 for each cell type). Error bars represent s.d. **D.** Quantification of tumor-initiating cell (TIC) frequency in control (Vector) and STAT3 Y640F-expressing KP MEFs by extreme limiting dilution assays (ELDA) in nude mice. **E.** Focus formation by KP MEFs expressing vector alone, STAT3 Y640F or double mutants of STAT3 Y640F with DNA binding domain (DBD) mutations, V463Δ and R382W, or phospho-inactivating Y705F. Results represent replicates from two independent experiments (n=4 for each cell type). Values correspond to mean ± s.d..

To determine whether specific Y705 phosphorylation and DNA binding are required for STAT3 Y640F to exert its suppressive activity, we generated three double mutants: a phosphorylation defective STAT3 Y640F/Y705F, and two DNA-binding domain (DBD) mutants; STAT3 Y640F/R382W and Y640F/V463Δ (Fig. S1E). Both R382W and V463Δ are recurrent STAT3 mutations observed in humans (30). A STAT3-responsive luciferase reporter assay was used to confirm that the DBD mutants are impaired in their ability to induce STAT3-mediated gene transcription (Fig. S1F). Notably, the DBD mutants of STAT3 Y640F lost the ability to attenuate KRAS-mediated MEF transformation despite their continuous pY705 phosphorylation (Figs. 2E, S1E). The phosphorylation defective STAT3 Y640F/Y705F double mutant was likewise impaired. We conclude that STAT3 Y640F-mediated inhibition of tumor development and KRAS-induced MEF transformation is dependent on STAT3 phosphorylation at Y705, DNA binding, and gene-specific transactivation.

To elucidate the means by which hyperactive STAT3 suppresses KP MEF transformation, we evaluated pathway activity by Western blot analysis, and transcription profile by whole exome RNA sequencing (RNA-seq). Western blot analyses of control and STAT3 Y640F-expressing MEFs showed unperturbed RAS/MAPK and PI3K/AKT signaling (as assessed by phosphorylated ERK1/2 and AKT1), suggesting that STAT3 does not directly alter these downstream KRAS effectors (Fig. S1B). RNA-seq analysis showed that expression of STAT3 Y640F in MEFs resulted in the differential expression of approximately 290 genes (p<0.05) compared to control cells. Gene ontology (GO) classification of biological processes showed an enrichment of pathways consistent with the role of STAT3 as a mediator of immunity and inflammatory response (Fig. S1G). Biological processes attenuated by hyperactive STAT3 included differentiation and tissue development, and pathways mediated by TGF-β family signaling (12, 13, 31). We therefore tested whether inactivation of the TGF-β pathway could inhibit KRAS-induced MEF transformation. Indeed, ablation of the TGF-β signaling components TGFBR2 or SMAD4 in KP MEFs using CRISPR/Cas9-mediated gene editing nearly eliminated foci formation and effects of STAT3 (Fig. SH, S1I). The results support the premise that hyperactive STAT3 interferes with KRAS-induced transformation through suppression of the TGF-β pathway. This prompted us to investigate the functional interaction of STAT3 and TGF-β/SMAD4 in epithelial carcinogenesis.

### STAT3 is a genetic modifier of EMT

A notable feature of KRAS mutant cancers, including those of the pancreas, colon, and lung, is that they tend to fall into two classes based on their canonical KRAS and TGF-β signaling; those that have a strong dependence on KRAS signaling (KRAS-dependent) or those that have less dependence on canonical KRAS signaling (KRAS-independent) (25, 26). KRAS-dependent tumors have been associated with an epithelial gene signature and morphology, whereas KRAS-independent tumors show enriched expression of mesenchymal genes. Since STAT3 and TGF-β have been shown to compete, cooperate, or antagonize each other in many other contexts (32–35), we investigated STAT3 as it relates to KRAS dependency. To that end, we used murine pancreatic ductal adenocarcinoma (PDAC)-derived cell lines bearing endogenous KRAS^G12D^ and TP53^R172H^ mutations (termed KPC) (17). CRISPR/Cas9 gene editing was used to ablate STAT3, SMAD4 or TGFBR2 expression in PDAC cells (Fig. S2A) and the behavior of these cells was tested alongside previously generated KRAS knockout cells (23). The cells were implanted orthotopically into the pancreata of nude mice, and animals were observed for latency of tumor formation and changes in tumor morphology.

Pancreatic tumors were detected within three weeks following implantation of 10^4^ parental control, STAT3 knockout (KO), SMAD4 KO, or TGFBR2 KO PDAC cells, and there was no statistical difference in tumor latency between the groups. However, there was considerable difference in tumor morphology. Tumors in the parental (intact) group were characterized by classical adenocarcinoma-like morphology with glandular structures (Fig. 3A). In comparison, KRAS KO tumors displayed a highly sarcomatoid morphology indicative of full EMT. Loss of STAT3 also induced a morphological change compatible with EMT, whereas overexpression of hyperactive STAT3 Y640F resulted in the sporadic co-occurrence of squamous and glandular differentiation. Loss of SMAD4 or TGFBR2 was associated with increased epithelial differentiation relative to controls. This is consistent with previous findings showing that the TGF-β pathway is a key regulator of tumor cell differentiation and malignant behavior, but not growth rate (36, 37).

**Figure 3.**
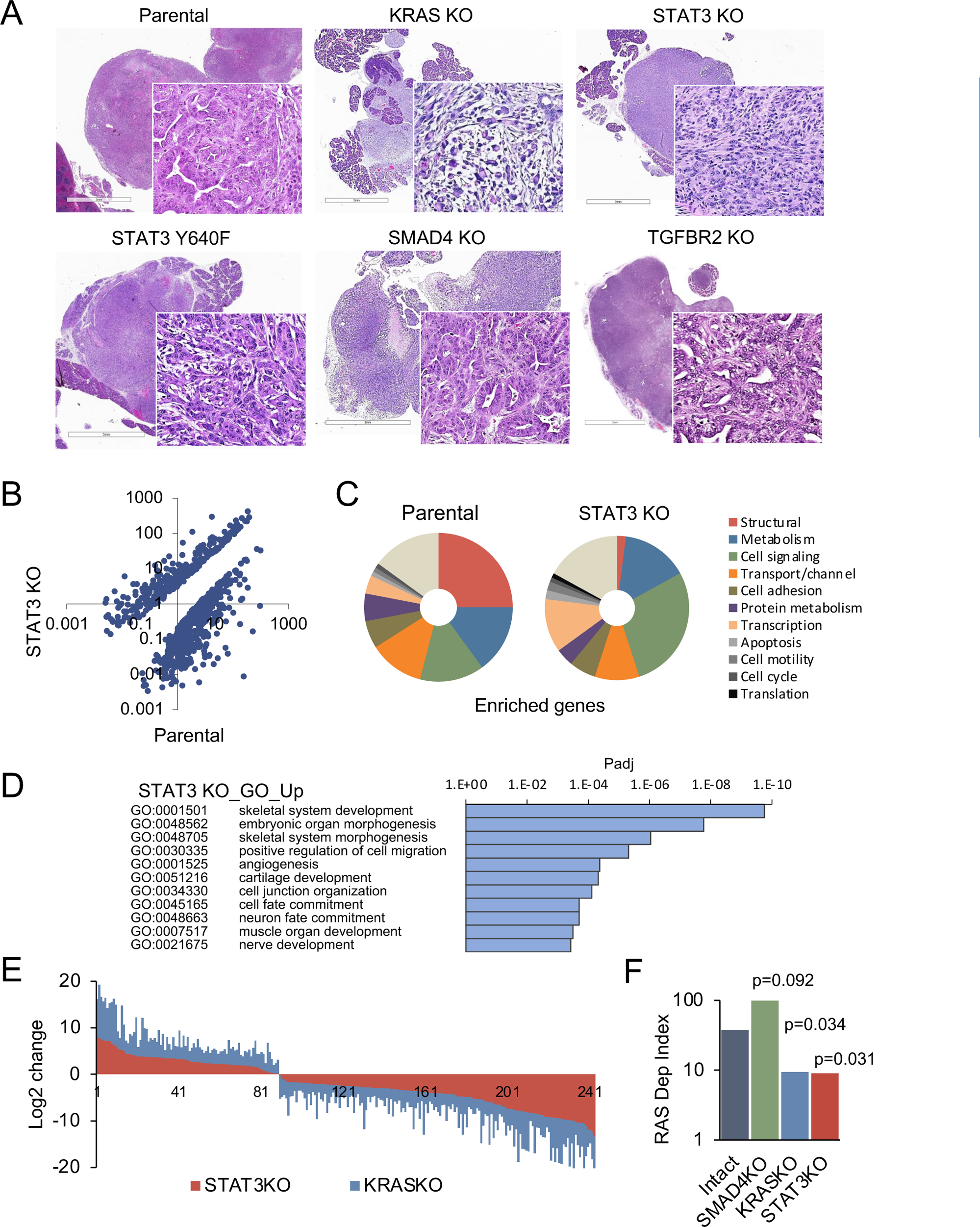
STAT3 is a genetic modifier of EMT. **A.** Representative H&E staining of pancreatic tumor sections derived from KPC cells of the indicated KRAS, STAT3, and SMAD4 genotypes. **B.** Scatter plot representing differentially expressed genes in parental (STAT3 intact) and STAT3 KO KPC cells. **C.** Pie charts representing gene expression profiles of the top 100 upregulated genes in parental control (STAT3 intact) and STAT3 KO KPC cells with percentages linked to specific cellular functions. **D.** GO enrichment analysis of the top 100 upregulated genes in STAT3 KO cells compared to STAT3 intact (control) cells. **E.** Analysis of overlapping differentially expressed genes (n=243) in STAT3 KO and KRAS KO KPC cell lines relative to parental control cells. **F.** Mouse RAS dependency index (RDI) of KPC cells of the indicated KRAS, STAT3, and SMAD4 genotypes.

We determined whether differences in tumor morphology were coordinate with changes in gene expression. Comparative RNA-seq analyses of parental KPC control and STAT3 KO cell lines revealed distinct transcriptional profiles that included more than 700 differentially expressed genes (Fig. 3B). The enriched genes in KPC parental cells included those corresponding to the major structural proteins in epithelial cells, such as cadherins, claudins, and tight junctions (Fig. 3C). In contrast, STAT3 KO cells were enriched in signatures of EMT and embryonic organ morphogenesis (Fig. 3D). The overall pattern of gene expression suggests that loss of STAT3 is associated with activation of partial rather than complete EMT, since epithelial markers (e.g. CDH1 and EpCAM) continue to be expressed, but mesenchymal markers (e.g. FN1 and various collagens) have been acquired (38, 39) (Figs. S2B). As TGF-β classically promotes EMT, we also focused on TGF-β family genes (40). Among these genes, STAT3 KO cells had a significant increase in TGFB1, TGFB3 and INHBA expression (Fig. S2C). The expression of EMT-activating transcription factors SNAI, TWIST and ZEB was not strongly affected, indicating that induction of EMT involves additional STAT3 dependent regulators. We did identify transcription factors, such as JUNB and SOX4, that associate with EMT (41). We computed EMT scores using gene expression values of epithelial (EPI) and mesenchymal (MES) genes. STAT3 and KRAS KO KPC cells displayed similar levels of EMT at the level of gene expression (Fig. S2D) (22). In contrast, SMAD4 KO cells displayed reduced expression of EMT-related genes, while genes involved in epithelial differentiation and RAS dependency were among the most upregulated (Fig. S2C).

Notably, comparative analysis between STAT3 and KRAS knockout KPC cells revealed approximately 250 STAT3 target genes (>30%) that were similarly up- or down-regulated (Pearson’s r=0.88, p<0.00001), suggesting that STAT3 loss partially phenocopies the effects of KRAS inactivation (Fig. 3E). GO classifications of the overlapping genes in KRAS and STAT3 knockouts included developmental processes and mesenchymal tissue remodeling (Figs. 3D, S2E). As a proof of concept for the functional connection of STAT3 to KRAS dependency, we used gene expression data to compute RAS dependency scores for parental and knockout cells. A mouse KRAS dependency signature (21 genes) was derived from single cell RNA-seq data of KRAS intact vs. KRAS knockout PDAC tumors and used for the analyses (Fig. S2F)(23). Results showed RAS dependency scores had a significant positive correlation with STAT3 expression and KRAS expression and a negative correlation with SMAD4 expression (Fig. 3E). Whole tumor RNA-seq analysis of STAT3 intact and knockout pancreatic tumors supported these findings (Fig. S2F). Together these data indicate that STAT3 is a genetic modifier that can regulate KRAS dependency and tumor development through counterposing EMT.

### STAT3 and SMAD4 play opposing roles in pancreatic tumorigenesis

The apparent antagonism between STAT3 and induction of EMT was of particular interest, as TGF-β-induced EMT appears to confer adaptive resistance to KRAS inhibition (25, 42). To test the relative importance of the STAT3 and TGF-β pathways to tumor morphology and functionality, we generated STAT3/SMAD4 and STAT3/TGFBR2 double knockout (DKO) cell lines. DKO cell lines formed pancreatic tumors in mice, but their histological features were distinct from SMAD4 or TGFBR2 single knockouts as they produced mixed epithelial/mesenchymal morphologies with cells expressing both E-cadherin and vimentin (Figs. 4A, 4B). SMAD4 or TGFBR2 intact tumors displayed features of EMT in the absence of STAT3, whereas SMAD4 or TGFBR2 KO tumors displayed a well-differentiated epithelial phenotype only in the presence of STAT3 (Figs. 3A, 4A). The data reinforce the notion that functional antagonism of STAT3 and TGF-β/SMAD4 controls PDAC development and KRAS dependency (Fig. 4C).

**Figure 4.**
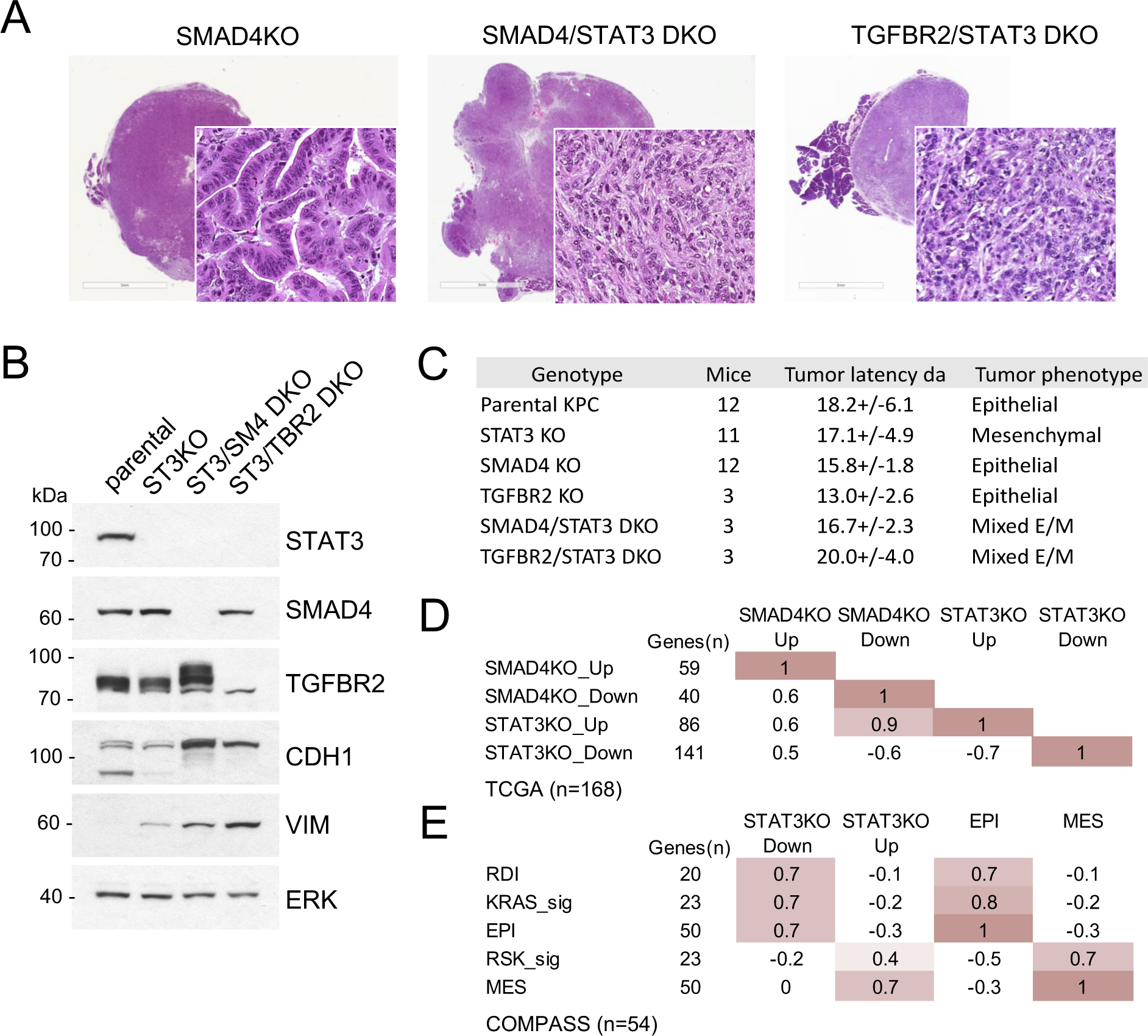
STAT3 and SMAD4 play opposing roles in pancreatic tumorigenesis. **A.** Representative H&E staining of pancreatic tumor sections derived from KPC cells of the indicated genotypes, SMAD4 KO (n=12), SMAD4/STAT3 DKO (n=3) and TGFBR2/STAT3 DKO (n=3). At least two independent clones were used. Scale bar = 100μm **B.** Western blot analysis of control (Intact), STAT3 KO (ST3KO), STAT3/SMAD4 double KO (ST3/SM4DKO), or STAT3/TGFBR2 double KO (ST3/TRB2DKO) KPC cells. Expression of STAT3, SMAD4, TGFBR2, CDH1 (E-cadherin), and VIM (Vimentin) is shown. ERK1/2 is a loading control. **C.** Summary of pancreatic tumor development in nude mice by KPC cells of the indicated STAT3, SMAD4, and TGFBR2 genotypes presented. **D.** Pearson correlation heatmap comparing gene expression of SMAD4 KO (SMAD4KO_up, down) and STAT3 KO (STAT3KO_up, down) KPC cells with human PDACs from TCGA (n=168). Signature scores were calculated using the top up- and down-regulated genes in STAT3 and SMAD4 KO KPC cells. The number of genes for each comparison is shown. **E.** Pearson correlation heatmap comparing gene expression signature scores of genes regulated in STAT3 KO KPC cells with pancreatic tumors from the COMPASS database (n=92) classified by RAS dependency index (RDI), KRAS-dependent signature (KRAS-sig), KRAS-independent signature (RSK-sig), epithelial (EPI) or mesenchymal (MES) gene signature score.

We analyzed human PDAC databases to determine if our STAT3 and SMAD4 gene expression signatures could be projected onto human tumors. SMAD4 and STAT3 signature scores were computed from the top up- or down-regulated genes in KPC cell lines (Fig. S3A). When aligned with human PDAC samples from the TCGA cohort (stage I/II tumors), STAT3 and SMAD4-regulated gene signatures demonstrated significant statistical correspondence, supporting the selective antagonism of STAT3 and SMAD4 (Pearson’s r>0.5, p<0.00001) (Fig. 4D). As SMAD4 is frequently deleted in PDAC, samples expressing only wild-type SMAD4 were manually curated from the TCGA cohort (n=112). Tumors were classified as either epithelial (EPI) or mesenchymal (MES) using previously characterized gene sets (Figs. S3B). Results demonstrate that STAT3-regulated gene expression is closely associated with epithelial differentiation, while SMAD4-regulated gene expression is strongly associated with EMT (Pearson’s r>0.7) (Fig. S3C). The association with KRAS dependency status was also revealing. PDAC tumor samples from the TCGA cohort were grouped as KRAS-dependent/KRAS type or KRAS-independent/RSK type based on previously derived KRAS dependency signatures (Fig. S3D)(25, 26). KRAS-dependent tumors showed enriched expression of epithelial genes (EPI) and reduced expression of mesenchymal genes (MES), whereas the KRAS-independent samples displayed the inverse, as previously reported (24–26). Importantly, the KPC STAT3 knockout gene signature (i.e. upregulated genes) co-segregated with human KRAS-independent/mesenchymal PDAC tumors, while the SMAD4 knockout gene signature (i.e. upregulated genes) co-segregated with human KRAS-dependent/epithelial tumors (Figs. S3C). A similar trend was observed for the PanCuRx Translational Research Initiative (COMPASS, stage IV PDAC) cohort mainly composed of liver metastases, as the STAT3-reliant gene signature was also enriched in epithelial and KRAS dependent samples (r>0.7, p<0.00001) (Fig. 4E). Liver is the main site of PDAC metastases, and pancreatic cancer metastases commonly display a stabilized epithelial phenotype (43, 44). Overall, results demonstrate that STAT3-regulated gene expression is closely associated with epithelial differentiation and RAS dependency, while SMAD4-regulated gene expression is strongly associated with EMT and RAS independence. These findings underscore our basic premise that there exists an epistatic antagonism between STAT3 and SMAD4, highlighting a new role for STAT3 as a genetic modifier in KRAS mutant cancer.

## Discussion

The findings presented in this study have significant implications in two main aspects. We provide evidence that the STAT3 transcription factor acts as a genetic modifier of EMT and KRAS dependency in a mouse model of pancreatic carcinogenesis. Our results demonstrate that neither persistent activation nor genetic ablation of STAT3 confers a selective growth advantage on tumor cells. Instead, STAT3 plays a crucial role in guiding the morphological and functional characteristics of tumors by inhibiting EMT and maintaining epithelial identity. Furthermore, our study uncovers an intriguing relationship between the SMAD4 and STAT3 transcription factors in PDAC. While the involvement of STAT3 in cancer has been predominantly associated with chronic inflammation and fibrosis (45, 46), our data underscore the significance of epistasis as a key factor that underlies functional antagonism between STAT3 and TGF-β/SMAD4 signaling. Our findings shed light on the regulatory mechanisms involved in cancer progression.

Oncogenic KRAS mutations are observed in approximately 90% of pancreatic cancers and less frequently in other cancer types. However, the role of KRAS in PDAC maintenance, once held to be nearly absolute, has shown limitations. A notable feature of KRAS mutant cancers, including those of the pancreas, colon, and lung, is that they can be either KRAS-dependent or KRAS-independent, based on the degree of their addiction to canonical KRAS signaling. The concept of KRAS dependency, originally introduced as a measure of oncogenic addiction following KRAS inactivation, has proven to be multifaceted. It integrates KRAS signaling outputs, with effector topologies, cooperating mutations, and environmental cues (23, 25, 26, 47). Cellular morphology (epithelial versus mesenchymal) appears to be one of the most noticeable manifestations of different degrees of KRAS dependency. Although SMAD4 classically promotes EMT and KRAS independence (25), approximately 50% of moderately to well-differentiated tumors in the TCGA cohort are, in fact, SMAD4 wild-type. This indicates that SMAD4 mutation alone is not sufficient to predispose cancer cells to a particular RAS phenotype. Our study emphasizes the mutual antagonism of SMAD4 and STAT3, and a fundamental role of STAT3 in the maintenance of epithelial cell identity. While SMAD4 wild-type tumors displayed features of EMT in the absence of STAT3, SMAD4 KO tumors displayed an epithelial phenotype only in the presence of STAT3. The results demonstrate that STAT3 and SMAD4 inversely contribute to oncogenic dependency. STAT3 sustains the KRAS dependent phenotype and tumor aggressiveness, with the potential for improved efficacy of anti-RAS drugs, whereas SMAD4 promotes KRAS independence at the expense of enhanced therapy resistance. Therefore, the epistatic relationship between SMAD4 and STAT3 has implications for tumor aggressiveness, metastatic propensity, and therapeutic resistance.

Genes involved in cancer (∼200 drivers validated to date) affect critical cellular processes, rendering them tumorigenic or tumor suppressive. The Cancer Dependency Map project sets out to model the genetic landscape of cancer in accordance with the oncogene addiction paradigm. By employing high throughput RNAi or CRISPR knockout screens across a multitude of cancer-derived cell lines, the goal is to broadly identify putative cellular dependencies for cancer therapy. While an important undertaking, not all cell lines align well with tumor samples in terms of mutations and gene expression profiles (48). STAT3 exemplifies this problem, as efforts to understand its role of STAT3 in cancer have resulted in conflicting reports that show either a positive or negative role in tumor development (4, 5). Large-scale analysis of patient-derived PDACs (n=84) and pancreatic cell lines from the Cancer Cell Line Encyclopedia (n=39) reveal low to medium levels of STAT3 Y705 phosphorylation (Fig. S3E). Endogenous phosphorylation/activation of STAT3 appears able to function within a narrow operating range in multiple solid tumor types.

Cancers are complex biological systems exhibiting inexplicable levels of intractability and unpredictability. For instance, even when challenged with the same lethal anticancer drugs used in vitro, cancers show remarkable resistance in vivo. Further complicating the analysis, there exist KRAS mutant cell lines whose survival and growth are no longer dependent on continued KRAS activity (23, 49, 50). This raises doubts regarding their eventual responsiveness to targeted anti-KRAS therapies. Here, we used genetic analyses to identify STAT3 as a relevant dependency in KRAS-driven cancer. While *in vitro* evidence indicates that STAT3 lacks classical driver properties, it nevertheless plays an essential role in cancer maintenance and epithelial-mesenchymal plasticity. This may explain the rarity of STAT3 GOF mutations in human cancers with mutant KRAS.

## Acknowledgements

This work was supported by NIH grant RO1CA236389 and the Carol M. Baldwin Breast Cancer Research Award to NCR, and the Catacosinos Cancer Research Award to OP. We wish to thank Fang Yuan Hao for his assistance, Jean Rooney in the Stony Brook University Division of Laboratory Animal Research for her technical assistance in mouse surgeries, and orthotopic implants, and Yan Ji from the Stony Brook University Histology Core.

## Competing interests

There are no competing interests

## Data availability

Human PDAC expression profiles from The Cancer Genome Atlas (TCGA) were downloaded as z-scores from cBioPortal (http://www.cbioportal.org), along with additional tumor and clinical annotations. PDAC datasets from the Amsterdam UMC (AUMC) (51) and PanCuRx Translational Research Initiative (COMPASS) (52) were used as described. The RNA-Seq data has been deposited in DRYAD https://datadryad.org/stash/share/k_W7vZ8uousKvh8tA3Ctmvxdp2-1yPNf8OiefhM00q8. The scRNA-seq data have been deposited in the GEO/SRA database under accession code GSE132582 [https://www.ncbi.nlm.nih.gov/geo/query/acc.cgi?acc=GSE132582]. Additional information is available from the authors on request.

## Supplemental Figure Legends

**Supplemental Figure 1.**
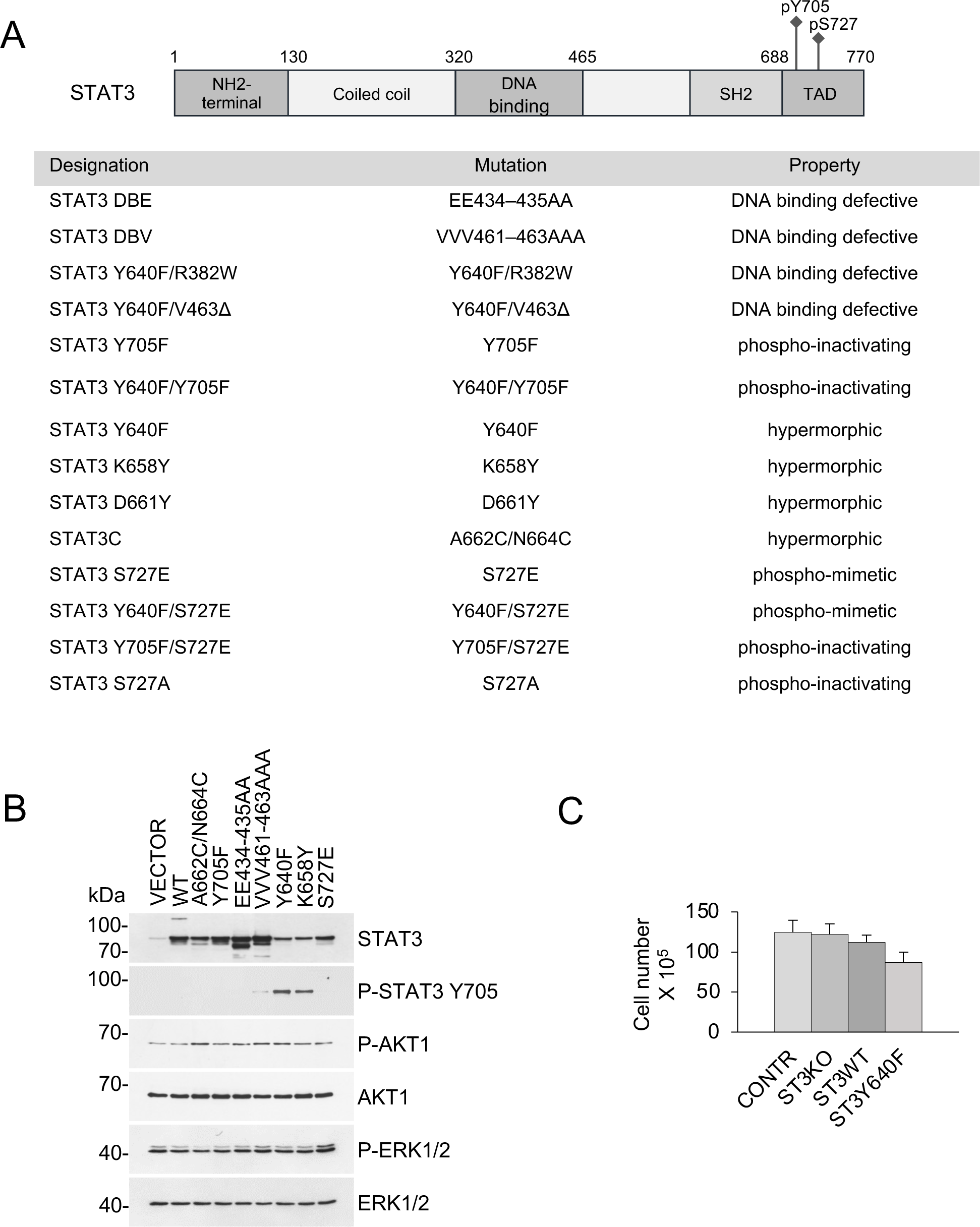
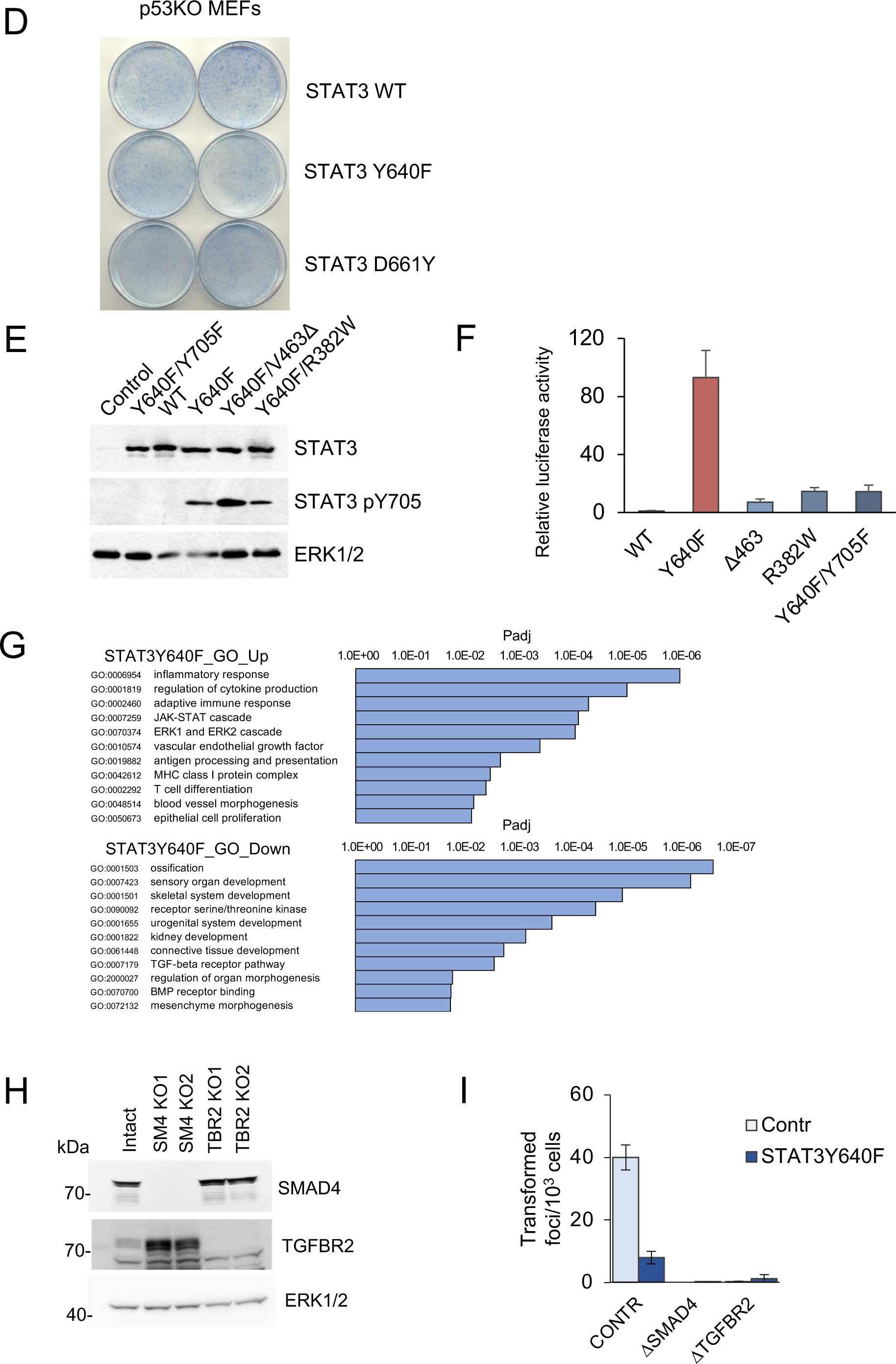
**A.** Schematic representation of STAT3 domain structure and the mutants used in this study. **B.** Western blot analysis of KP MEFs harboring STAT3 mutations reveals unperturbed PI3K/AKT and MAPK/ERK signaling. **C.** Loss of STAT3 does not affect growth and viability of KRAS^G12D^ p53KO MEFs. The cells were cultured for two weeks in 2D adherent cultures. Cumulative cell numbers are shown. Values correspond to mean ± s.d. **D.** Focus formation assays of p53null MEFs overexpressing STAT3 WT, STAT3 Y640F or STAT3 D661Y. **E.** Western blot analyses of total STAT3 and Y705-phosphorylated STAT3 in KP MEFs expressing vector alone, and single and double mutations of STAT3 as noted. ERK1/2 is a loading control. **F.** Transactivation by STAT3 mutants of a STAT3-responsive promoter driving a luciferase reporter. Hep3B cells were co-transfected with the p3XGAS-Hsp70-Luc reporter plasmid and mutant STAT3 alleles as noted and stimulated with 20 ng/ml human IL-6 for 24 hours (n=3 for each cell type). Relative luciferase values are presented as mean ± s.d. **G.** GO enrichment of differentially expressed genes in KP MEF cells expressing STAT3 Y640F compared to control KP cells provided by Novogene Corp. from RNASeq data. **H.** Western blot analysis of SMAD4KO and TGFBR2KO KP MEFs. **I.** Results of focus formation assays by control KP MEFs, and MEFs ablated for SMAD4 and TGFBR2 expression with or without expression of STAT3Y640F as indicated (n>6 for each). Values correspond to average and s.d.

**Supplemental Figure 2.**
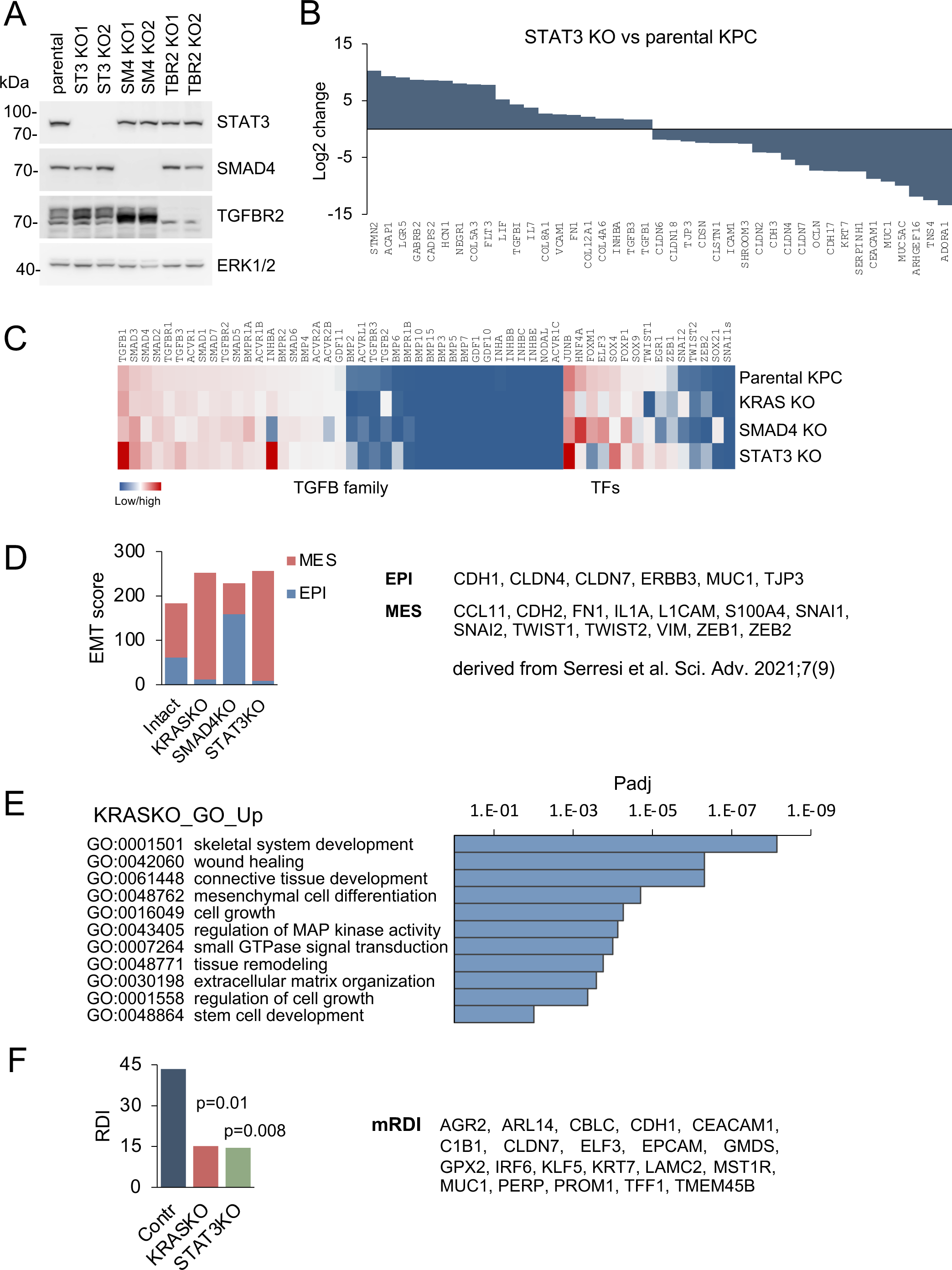
**A.** Western blots of parental KPC pancreatic ductal adenocarcinoma cells, and two isogenic knockout clones for STAT3, SMAD4 or TGFBR2 generated by CRISPR/Cas9 gene editing. ERK1/2 is a loading control. **B.** Top up- and down-regulated subset of genes in STAT3 KO KPC cells compared to parental control cells. **C.** Heatmaps of differentially expressed genes in KPC parental cells and the indicated KRAS, STAT3, and SMAD4 knockout genotypes. TGF-β super family genes and transcription factors (TFs) are shown. **D.** EMT scores of KPC cells of the indicated KRAS, STAT3, and SMAD4 genotypes based on epidermal (EPI) and mesenchymal (MES) genes derived from Serresi et al. Sci. Adv. 2021;7(9) shown to right. **E.** GO enrichment analysis of top up-regulated genes in KRAS KO cells compared to parental KPC cells. The normalized FDR (Padj) was provided by Novogene Corp. from RNASeq data. **F.** RAS dependency indices of KPC pancreatic whole tumors based on bulk RNA-seq of tumors formed from parental KPC (Contr), KRAS KO or STAT3 KO cells. Murine RAS dependency index (mRDI) was derived from Ischenko et al. Nature Communications, 2021;12(1):1482.

**Supplemental Figure 3.**
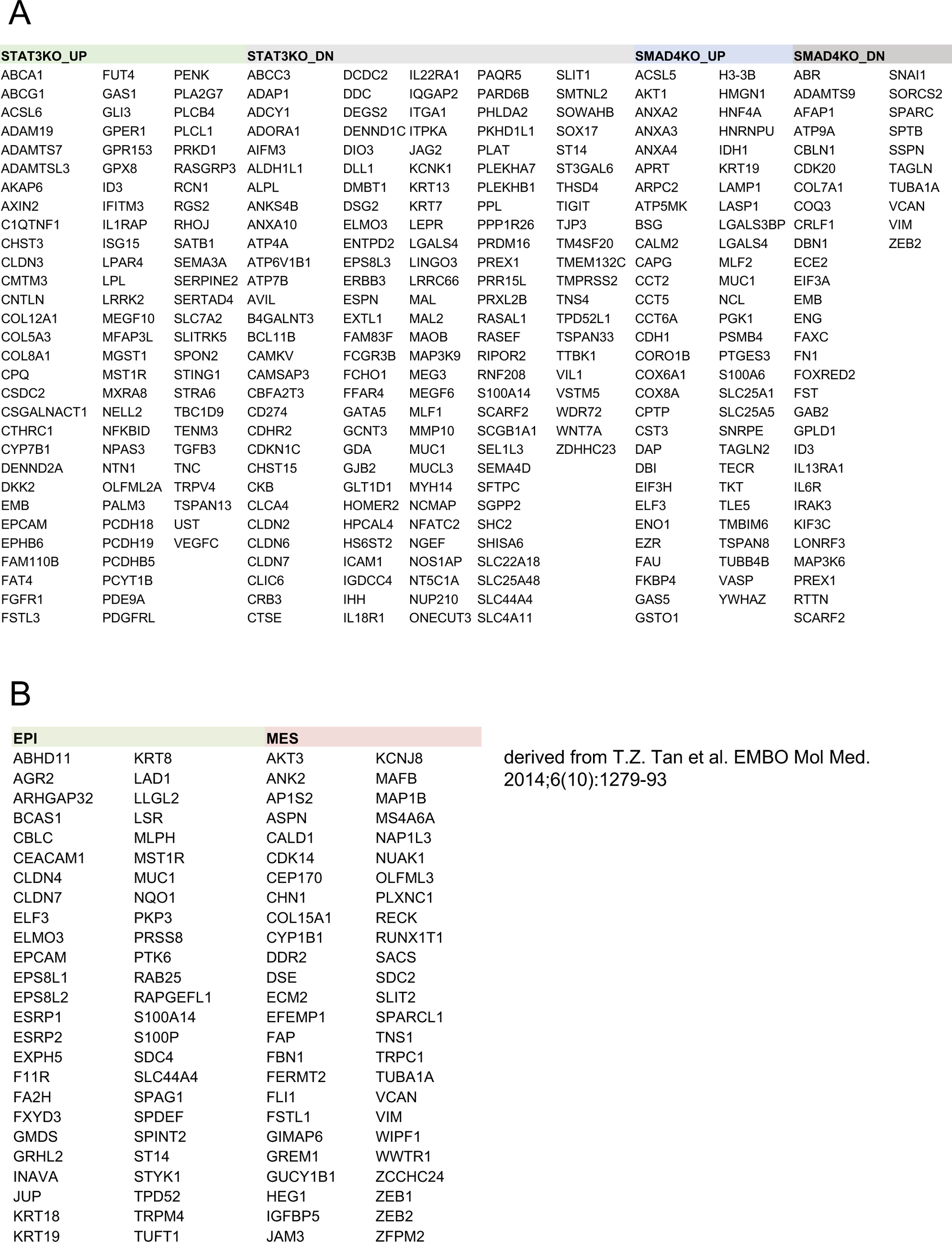
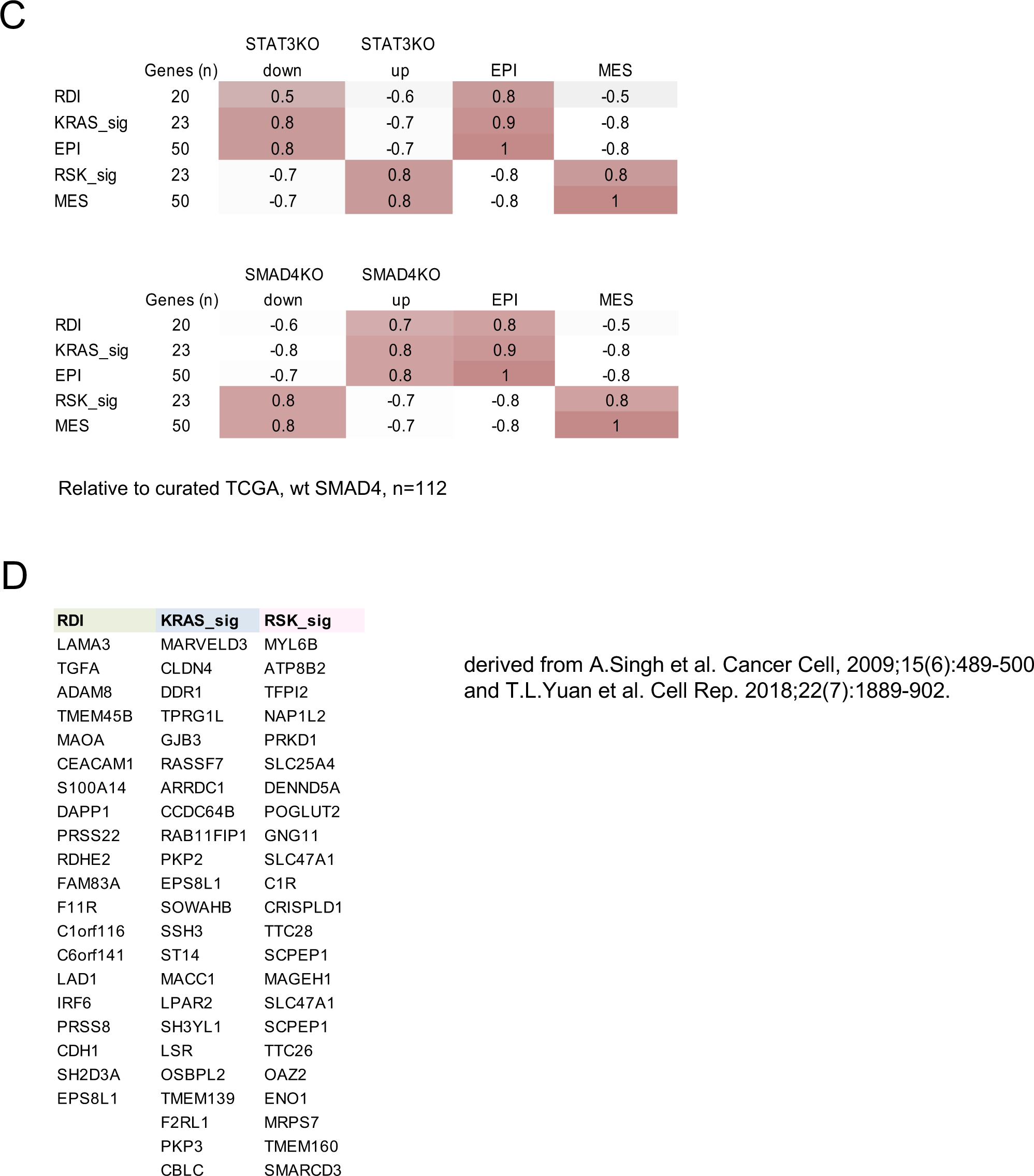
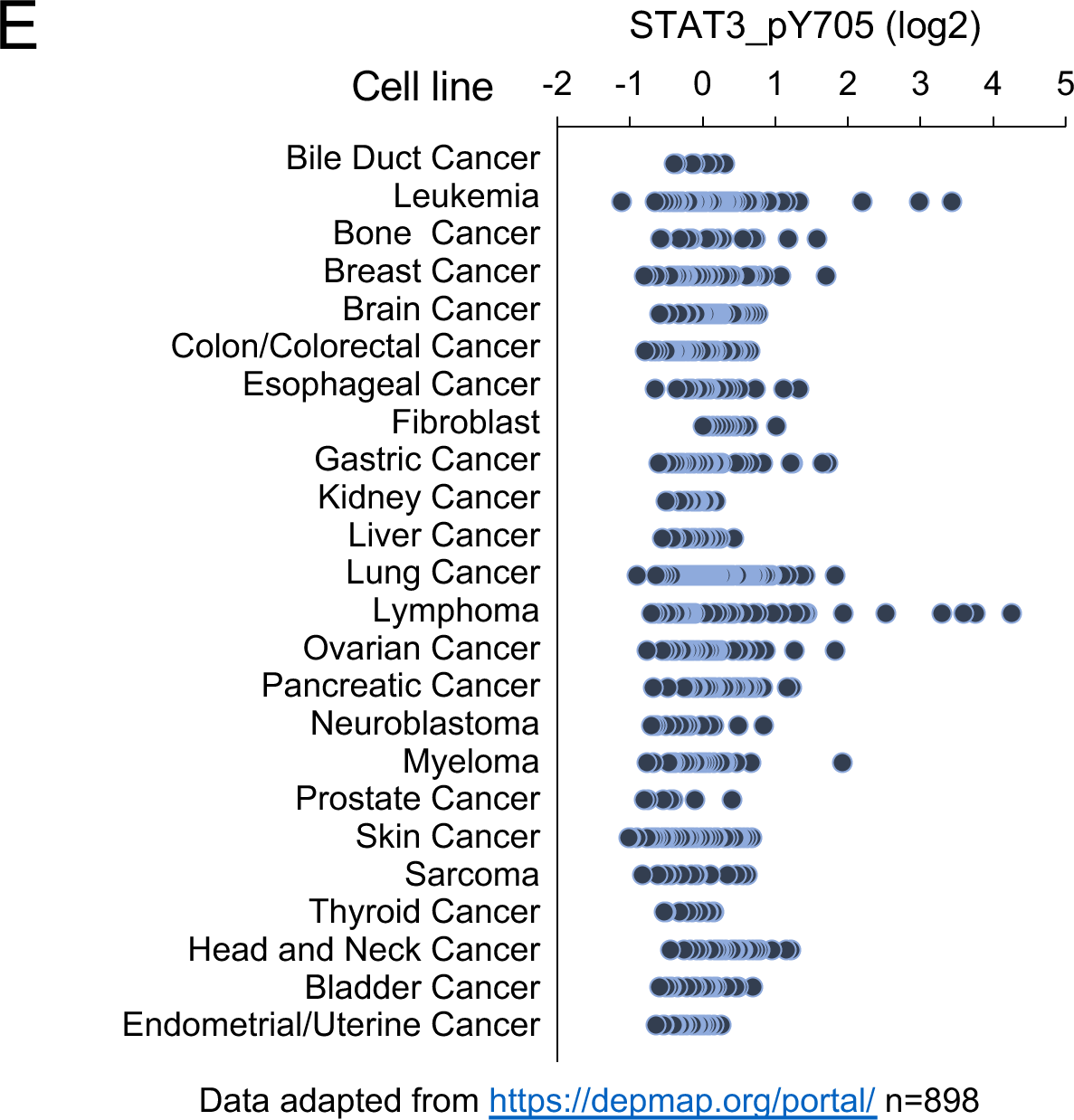
**A.** Top up- or down-regulated genes in murine KPC STAT3 KO or SMAD4 KO cells derived from RNA-Seq. **B.** Signature genes used to designate TCGA and COMPASS PDAC individual tumors as epithelial (EPI) or mesenchymal (MES) derived from T.Z. Tan et al. EMBO Mol Med. 2014;6(10):1279-93. **C.** Pearson correlation coefficients of STAT3 KO gene signature scores (top) or SMAD4 KO gene signature scores (bottom) with curated human pancreatic tumor samples of expressing wild-type SMAD4 from TCGA (n=112) classified by Ras dependency index (RDI), KRAS-dependent signature (KRAS-sig), KRAS-independent signature (RSK-sig), epithelial (EPI) or mesenchymal (MES). The number of genes for each comparison is shown. **D.** Genes used to define a RAS Dependency Index (RDI) derived from A.Singh et al. Cancer Cell, 2009;15(6):489-500, and genes used to define a KRAS dependent signature (KRAS_sig) or an RSK dependent signature (RSK_sig) derived from T.L.Yuan et al. Cell Rep. 2018;22(7):1889-902. **E.** STAT3 Y705 phosphorylation levels in cell lines from the Cancer Cell Line Encyclopedia (n=898).

## Notes

https://datadryad.org/stash/share/k_W7vZ8uousKvh8tA3Ctmvxdp2-1yPNf8OiefhM00q8

https://www.ncbi.nlm.nih.gov/geo/query/acc.cgi?acc=GSE132582

### Competing Interest Statement

The authors have declared no competing interest.

